# Differential Regulation of Large-scale Chromosome Conformations in Osteoblasts and Osteosarcoma

**DOI:** 10.1101/2024.11.01.621571

**Authors:** Madhoolika Bisht, Yu-Chieh Chung, Siou-Luan He, Sydney Willey, Benjamin D. Sunkel, Meng Wang, Benjamin Z. Stanton, Li-Chun Tu

**Author notes:** Present address: Michigan Neuroscience Institute, University of Michigan, Ann Arbor, MI 48109. These authors contributed equally to the work (Madhoolika Bisht, Yu-Chieh Chung).

## Abstract

Correct chromosome organization in the cell nucleus is essential for genome function. However, dynamics and regulations of large-scale chromosomal conformations beyond single compartments at larger than ten megabases *in vivo* in single cells remain largely unknown. Here we use CRISPR-Sirius, a high-resolution and high-sensitivity real-time imaging technique, to directly visualize distinctions in large-scale chromosomal conformations between live osteosarcoma (OS) cells and osteoblasts, suggesting extensive chromatin reorganization during cell transformation. A surprising discovery is that chromosome 19 long arm is primarily extended in osteoblasts and maintained by H3K27me3. Extended chromosome conformation has been reported in fly and mouse but not in human cells yet. However, in OS cells, chromosome 19 primarily folded into collapsed conformations, which reshape in minutes and are regulated by the chromosome architectural proteins CTCF and cohesin in the presence of H3K27ac. Changes in chromosome conformations by knocking down the cohesin subunit RAD21 resulted in altered gene expression, including proto-oncogenes. Transcription inhibition by a small molecule inhibitor did not have detectable effects on large-scale chromosome conformation, suggesting that local transcription events have limited effects on large-scale chromosomal architecture. Our results provide unique insights into the complex regulatory mechanisms of endogenous large-scale chromosome organization in normal and transformed osteogenic tissues.

**One-sentence summary:** Regulations of chromosome territory organization in normal and cancer bone cells visualized by CRISPR-based live-cell imaging.

## INTRODUCTION

Three-dimensional (3D) chromatin architecture is dynamically and hierarchically organized in the cell nucleus(1). Chromatin compaction controls DNA accessibility and is crucial for cellular processes, such as transcription and replication(1, 2). The fundamental unit of DNA compaction is the nucleosome, in which DNA wraps around the histone octamers (3). However, evidence for decades has shown that despite their utility in modulating DNA conformational states, nucleosomes are highly dynamic, and unwrap through entropically driven processes in addition to ATP-dependent remodeling (4). Distances between nucleosomes are also regulated by linker histones and posttranslational modifications (PTMs) on histone tails, such as methylation and acetylation(5), which can serve to alter local electrostatic environments or alter nucleosome engagement by histone reader domains (6). Histone PTMs also regulate gene expression by recruiting transcriptional cofactors, including remodelers and silencers (7). Dysregulation of histone PTMs has been reported in multiple cancers(8, 9).

Specific patterns of histone methylation and acetylation can alter chromatin compaction, which has been used to characterize chromatin states. Methylation of histone H3 at lysine-4 (H3K4me3) and acetylation of histone H3 at lysine-27 (H3K27ac) are known euchromatin marks that are associated with open chromatin (10). Both H3K4me3 and H3K27ac are associated with essential gene expression, and the combination of these histone modifications occurs at active promoters and enhancers (11). Conversely, heterochromatin marks are often found in densely packed chromatin and are involved in gene repression. Methylation of histone H3 at lysine-9 (H3K9me2/3) and lysine-27 (H3K27me3) are the most well-studied heterochromatin marks, each associated with heritability through read-to-write mechanisms (6). Each histone modification is added, removed, and read by specific enzymes. In the case of H3K27me3, a methyl group is added by the polycomb repressive complexes (PRC)(12) and removed by the demethylases UTX and JMJD3(13).

Although chromatin can intrinsically loop at dozens of kilobases, chromosome conformation capture-based techniques have revealed topologically associating domains (TADs)(14, 15), the chromatin loops regulated by the chromatin architectural proteins, CCCTC binding factor (CTCF) and cohesin(16, 17). There is evidence that TADs, with a median size of 360 kb in length, extrude through cohesin and are anchored by two convergent CTCF molecules at loop boundaries (18, 19). At sub-megabase scale, clusters of loops are partitioned into compartment A and B. Compartment A is gene rich and often marked by euchromatin marks, whereas compartment B is often gene poor and commonly marked by heterochromatin marks (20). Compartments are thought to be dynamic and heterogeneous among cells, therefore making them hard to be modeled(19). To date, a few compartments have been visualized in fixed cells of specific types but their biological importance is not clear (21, 22). Thus, segregation and conversion between compartments in human cells remain largely enigmatic. Each interphase chromosome contains several compartments and occupies a distinct 3D nuclear space known as chromosome territory, which limits interchromosomal interactions, thereby maintaining genome integrity(23).

Aberrant 3D genome organization has been found in cancer, and key studies have revealed that genome structure can serve as an oncogenic driver. For instance, altered 3D genome architecture in prostate cancer is correlated with deletion of tumor suppressor gene *TP53*(*24*). Similarly, the promoter-enhancer loops regulated by the presence of estrogen receptors (ERα) are altered in breast cancer(25). Analysis of somatic mutations from ∼42 different human cancer types showed that changes in the cancer genome largely occurred at the boundaries of TADs, indicating a strong association with changes in spatial genome organization(26, 27).

Advances in high-throughput sequencing and fluorescence in-situ hybridization imaging techniques have greatly built our knowledge on 3D genome organization in high resolution. However, these techniques require samples to be fixed, limiting the understanding of genome organization beyond size and shape. Whether euchromatin forms a static and open structure in all cell types is unclear(28). Additionally, it remains a key question in the field, whether different 3D genome architectures are instructive or responsive in cell fate decisions and how different regulatory mechanisms orchestrate to make epigenetic changes that affect gene expression across cell types, with and without a disease. Previous studies have revealed that gene expression drives changes in local chromatin organization(29–31). Chromatin topologies in mice are cell-type dependent, suggesting an integration of lineage specification and architectural signature(32). Despite being highly challenging, temporal evolution of chromosome organization at single chromosome level in live single cells helps to elucidate the interplay of genome function and structure in detail, which is not available in experiments using fixed cells. Human genome organization in real time at chromosome territory scale (>10 Mb) containing dozens, if not hundreds, of genes is less known.

Live cell imaging techniques such as CRISPRainbow and CRISPR-Sirius(33–35) have shown its power to provide significant insights into dynamic chromatin organization. Early live-cell imaging techniques were limited to specific regions, such as telomeres and centromeres, by their DNA-binding proteins (36). Whole genome labeling, such as fluorescent histones, have revealed dynamics of chromatin architecture(37). However, these techniques do not have gene and chromosome specificity, thus limiting questions to be answered. Insertion of bacterial lac operon allows local imaging at specific genomic loci but potential effects from foreign DNA bound by exogenous proteins, such as interactions with PML bodies, were reported (38, 39). CRISPR imaging has been used to study chromatin dynamics on well-characterized endogenous genomic sequences, including telomeres, pericentromeric regions, active versus silence genes, and TADs’ boundaries(40–42). The high precision of CRISPR had been used to visualize aneuploidy and chromosome translocation in cancer cells (43). In this study, we utilized CRISPR-Sirius to investigate large-scale chromatin conformation at > 10 Mb scale in osteosarcoma (OS) and osteoblast cells. We uncovered striking differences in 3D conformations of chromosome 19 territory between osteoblasts and OS. As of today, the extended chromosome conformations were only reported in fly and mouse, but not in human cells yet (44). In this work, we uncovered the stability of extended chromosome territories in live human cells which has not been characterized before. We found that inhibition of transcription alone is not sufficient to reorganize the *in vivo* large-scale chromosomal conformations, but changes in chromosomal conformations are associated with altered gene expression. Distinct chromatin regulators with little or no cross-talk are used in controlling the osteogenic 3D chromosome conformations. Our and others’ previous work revealed heterogeneous organization between homologs (41, 45). However, whether different chromosome territory organizations are results of temporal morphological dynamics or cell-to-cell variety is unclear. Here we show that human chromatin organization at > 10 Mb scale reshapes in minutes in living cells, which likely represents a general phenomenon of gene-rich chromosomes in higher eukaryotic genomes *in vivo*.

## RESULTS

### Chromosome 19 long-arm conformations and gene profiles are distinct in osteoblasts (OB) and osteosarcoma (OS)

We used CRISPR-Sirius to visualize and investigate large-scale chromosome conformations in living cells(34). In CRISPR-Sirius system, the crRNA and tracrRNA were genetically fused to a single guide RNA (sgRNA) scaffold engineered with eight PP7 aptamers (8xPP7) (Figure 1A). The fluorescent PP7 coat protein (PCP) was made by fusing with green fluorescent protein (PCP-GFP). CRISPR-Sirius sgRNA is thermostable in the human nucleus. The nuclease-dead CRISPR-associated protein 9 (dCas9) has two point mutations at D10A and H840A to deactivate its endonuclease activity(46). These plasmids have been reported in previous work(33, 34).

**Figure 1.**
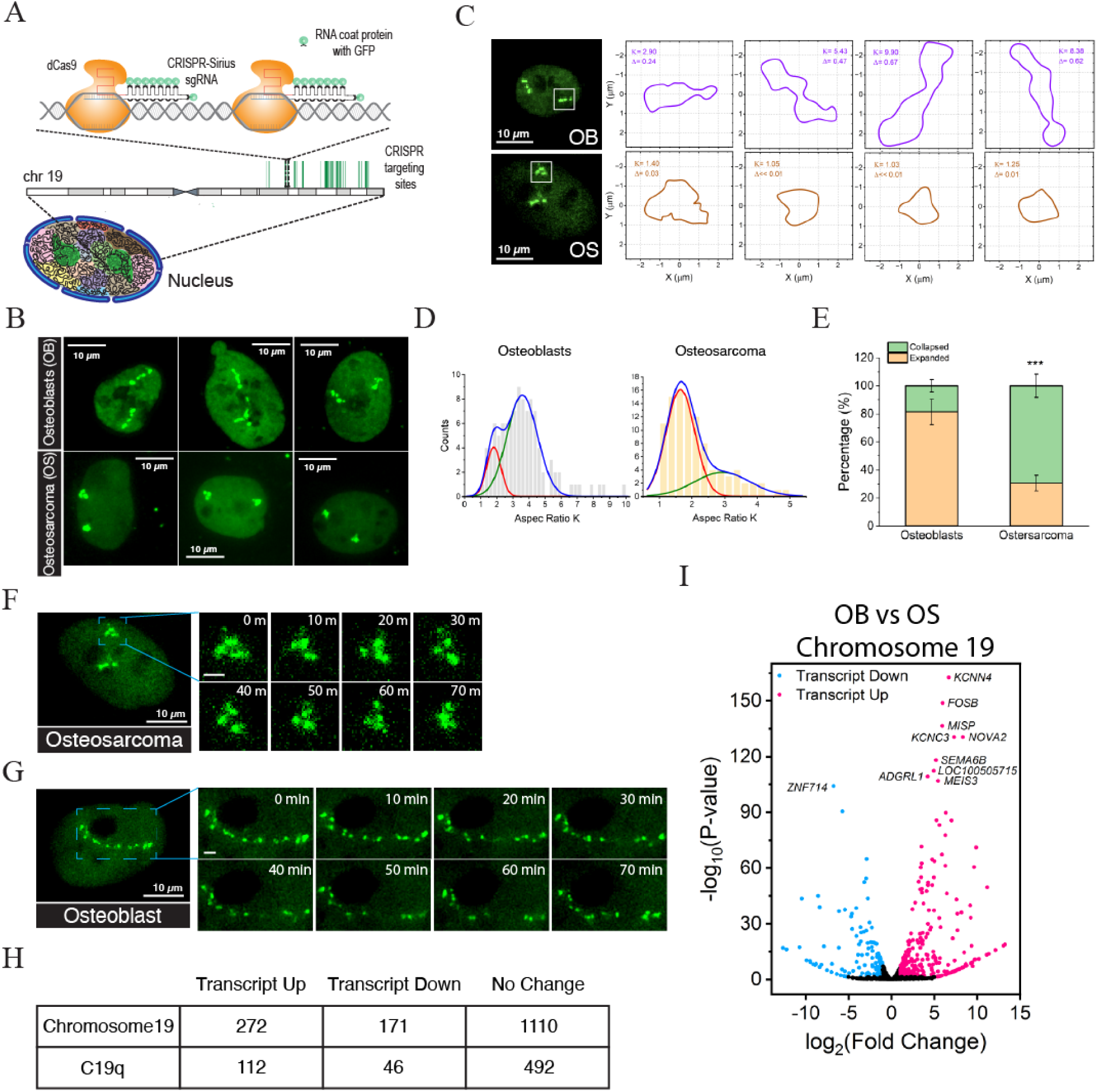
Distinct chromosome conformations in osteoblasts and osteosarcoma cells. (A) A schematic of CRISPR-Sirius. The CRISPR-Sirius single guide RNA (sgRNA) is engineered with an octet of PP7 hairpins (8XPP7). PP7 is labeled by binding to its coat protein (PCP) fused with the green fluorescent protein (GFP). The sgRNA targeted about 836 repeats spanning 17 Mb region on chromosome 19 q arm (C19q), which allows the visualization of chromosome 19 territory. (B) Images of C19q conformation in osteoblasts (OB, hFOB1.19) and osteosarcoma (OS, U2OS) cells. Osteoblasts have extended C19q conformations, whereas osteosarcoma cells demonstrated collapsed C19q conformations. (C) Schematic of C19q conformation quantitative analysis. The spline curves from C19q images in Fig 1B and 1C were used to calculate gyration tensors that give rise to the asphericity (Δ) and aspect ratio (K) of C19q conformations in osteoblasts (upper row) and osteosarcoma cells (lower row). (D) Gaussian bimodal fitting of aspect ratio (K) distribution of C19q conformations for osteoblast (N_C19q_= 104) and osteosarcoma cells (N_C19q_=100). The blue lines represent fittings with two Gaussian peaks for whole distributions. Red and green lines represent the individual distributions for collapsed and extended conformations, respectively. (E) The percentages of collapsed and extended conformations from gaussian bimodal fittings. Histogram shows that osteosarcoma cells (N_C19q_=100) (***, p < 0.001) have significantly fewer C19q in the extended conformation as compared to those in osteoblasts (N_C19q_= 104). (F) Time lapse of C19q over 70 minutes in osteosarcoma under physiological conditions (37°C, 5% CO_2_ and humidity)(G) Time lapse of C19q over 70 minutes in osteoblasts under physiological conditions (37°C, 5% CO_2_ and humidity). The m denotes minutes. Bar in the enlarged inset, 2 µm. Each Z-stack was 3D deconvoluted and projected using maximal intensity. (H) Numbers of genes up– and down-regulated in chromosome 19 and in C19q, respectively. (I) Volcano plot of RNA-seq comparing differential expression of chromosome 19 genes in OB and OS. The significance test in (E) was calculated using Fisher’s exact test. All experiments were repeated at least three times.

We selected human chromosome 19 (chr19) for this study because (1) it is a gene-rich chromosome. The gene density of chr19 is more than double the genome wide average(47), and (2) mutations in chr19 have extensive correlation with cancer progression and poor prognosis(48–53). To determine the conformation of chr19, we used CRISPR-Sirius to target ∼ 836 repetitive sequences on the long arm (C19q) spanning across an ∼17 Mb region between 19q13.2 and 19q13.42 (Figure 1A)(54). The C19q region was fluorescently labeled in hFOB1.19 cells, hereafter referred to as osteoblasts (OB), and U2OS cells, hereafter referred to as osteosarcoma (OS) cells, using lentiviral transductions to generate stable cell lines (Figure 1B). The region targeted by C19q has been verified in U2OS cells through previous studies using FISH oligopaint(54).

Upon acquiring C19q conformations using 3D imaging, we observed different preferential conformations, extended and collapsed, in osteoblasts and OS cells, respectively (Figure 1B). To quantify C19q conformations, we calculate C19q asphericity (Δ) which has been used to characterize the conformation of polymers (55, 56) and is given by the eigenvalues of gyration tensor calculated from the spline curve of C19q (Fig 1C) (Materials and Methods). Asphericity is a dimensionless quantity between zero and one. Δ=0 represents a circular shape while Δ=1 represents a fully extended shape. The smaller value, the rounder conformation. To obtain a more geometric picture, it is instructive to calculate another dimensionless quantity, aspect ratio K of the principal axes of gyration tensor. When K=1, the conformation is round and the bigger K, the more extended conformation. We then found that the distribution of K values can be fitted into Gaussian bimodal distributions (Figure 1D). The mean of K values for collapsed and extended C19q of OS are 1.65 and 2.93, respectively, while osteoblasts have mean K=1.82 and K=3.58 for collapsed and extended conformations of C19q, respectively. In osteoblasts, approximately 81% of C19q conformations were extended. In contrast, only ∼31% of OS cells displayed extended C19q conformations (Figure 1E, total number of cells, n≥100, \*\*\**p*<0.001). C19q was present in predominantly collapsed conformations in OS cells (69%) (Figure 1E). To examine the conformational stability, we tracked temporal conformational changes by recording 3D time-lapse images at 10-minute intervals. We found that the collapsed C19q in OS cells continued to collapse but underwent local reshaping in 10 minutes (Movie S1 and Figure 1F). Similarly, the extended C19q in osteoblasts remains extended for 70 minutes (Movie S2 and Figure 1G). To examine whether the different C19q conformations result from different cell-cycle stages, we examined C19q conformations in synchronized osteoblasts and OS cells (Figure S1). We synchronized both cell lines to G1 phase with a double thymidine block followed by nocodazole treatment(41, 57, 58). C19q conformations were captured 6 hours after nocodazole release, which corresponds to the mid-late G1 phase. Similar results were obtained to those in asynchronized cells, suggesting that the differences in C19q conformations in the two cell lines were not due to differences in the stages of the cell cycle but indicate the ability of 3D chromatin architecture to reorganize in disease conditions such as OS.

Next we asked whether the distinct C19q conformations in OS and OB are correlated with different gene expression profiles. We examined the total transcripts coded in chr19 and within C19q comparing RNA-seq analysis of the two lines (Figure 1H and 1I). About 443 of chr19 genes (29%) were differentially expressed in OS cells, and 158 (36%) of them are within the C19q region. Whole genome gene set enrichment analysis (GSEA) in oncology showed that bone cancer related genes are dysregulated in OS, such as immune response, quality control of damaged cells, osteogenic tissue identities, and abnormal tissue metabolite concentrations (Figure 4I). OS cells have reduced response to virus and radiation, signs of weakened immune system and radiotherapy resistance. The reduced osteoclast development, on the other hand, weakens the ability to remove old and damaged bone tissue. OS cells also have reduced bone cell characteristics by depleting genes related to bone development, bone mineralization, and regulation of cell adhesion, which might correlate to the weakened bones in patients with osteosarcoma.

### Transcription inhibition alone has undetectable effects on C19q chromosome conformations

Previous studies showed that transcription activation leads to an extension of the local chromatin fiber of the *HSP70* gene on bacterial artificial chromosomes in Chinese hamster ovary (CHO) cells, which is rescued by the transcription inhibitor, 5,6-dichloro-β-D-ribofuranosyl-benzimidazole (DRB) (29). To evaluate the effects of transcription on large-scale chromosome conformation, we inhibited transcription using DRB, which halts RNA Pol II elongation by inhibiting the CTD kinase(59). By measuring newly synthesized RNA using 5-ethynyl uridine (EU), we verified that transcription was successfully inhibited in osteoblasts and OS cells within 2 hours (Figure S2). We recorded changes in chromosome conformation for 2 hours in osteoblasts and OS cells during DRB treatment (Figure 2A and 2B). Surprisingly, transcription inhibition did not have detectable effects on C19q conformational states in either cell type (Figure 2C and 2D, total number of cells, n≥ 50, p>0.05), indicating that transcription does not regulate large-scale chromatin conformations. Similar results have been reported in HT-1080 cells during early G1 stage (60). Transcription inhibition by Actinomycin D did not affect the volume, surface area, and sphericity of human chromosome 11 territories. As a control, we examined the transcript levels of three active genes, NECTIN2 (42.9 kb), TOMM40 (12.4 kb), and TNNT1 (16.5 kb) in OS cells, targeted by C19q-sgRNA 8, 1, and 16 times, respectively. The active genes targeted by CRISPR-Sirius have no effects on the levels of transcripts, as demonstrated by the RT-PCR (Figure 2E).

**Figure 2.**
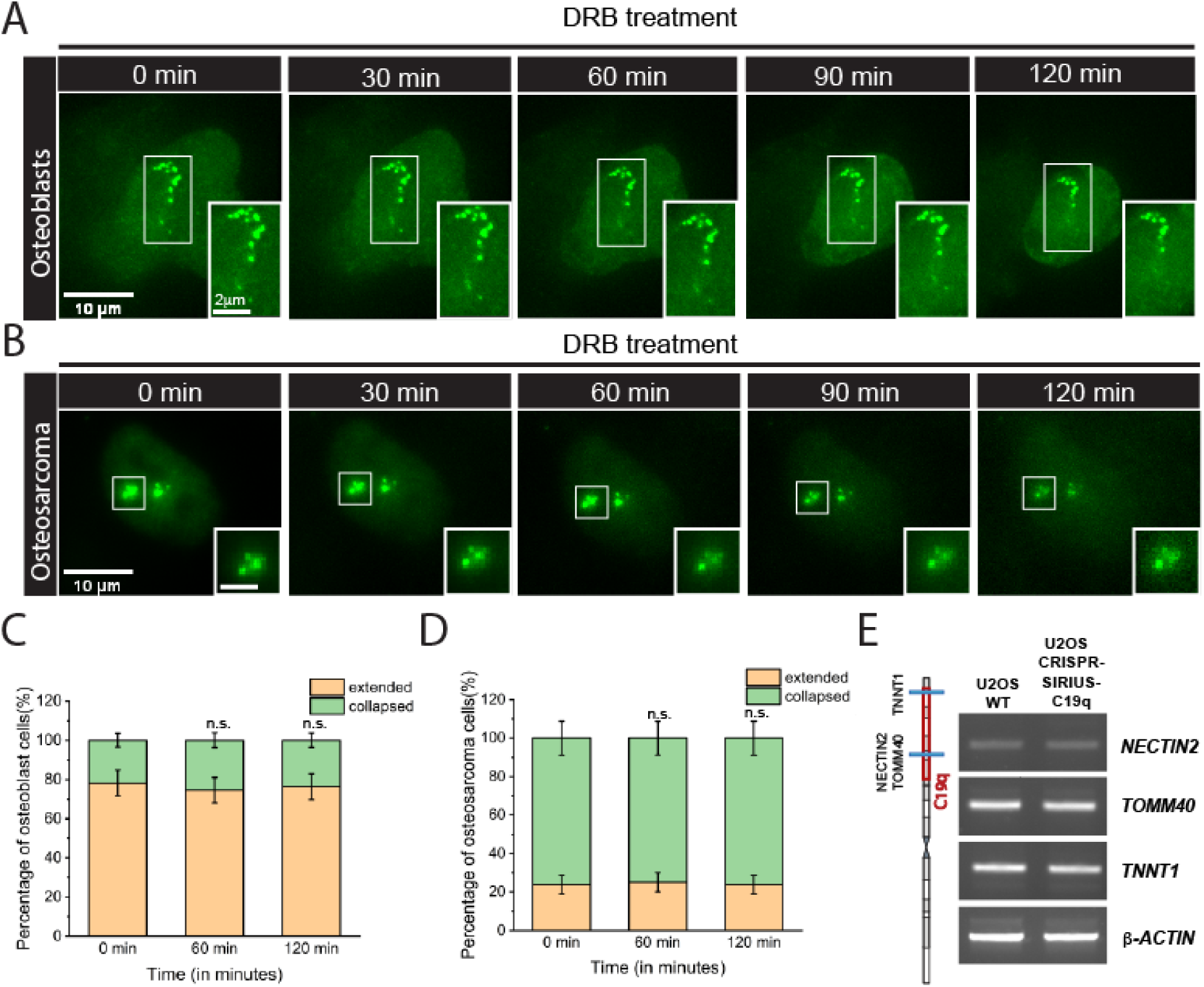
Transcription inhibition has no detectable effects on large-scale chromosome conformation. (A) Time-lapse images of chromosome conformation under DRB treatment for 120 minutes in osteoblast cells. Transcription inhibition by DRB treatment does not change the extended chromosome conformation. (B) Time-lapse images of chromosome conformation under DRB treatment for 120 minutes in osteosarcoma cells. Transcription inhibition does not affect the collapsed chromosome conformation in osteosarcoma cells. (C) Percentage of osteoblast cells with extended and collapsed C19q conformation were measured at 60 minutes (N_C19q_=55, p= 0.8228) (n.s., non-significant) and 120 minutes (N_C19q_=55, p-value = ∼1) (n.s, non-significant). (D) Percentage of osteosarcoma cells with extended and collapsed C19q conformation were measured at 60 minutes (N_C19q_=80, p-value = ∼1) and 120 minutes (N_C19q_=80, p-value = ∼1). n.s., Non-Significant. The significance test was calculated by using Fisher’s exact test. (E) RT-PCR of gene transcripts in C19q in OS cells with and without CRISPR-Sirius C19q-sgRNA targeting. Positive cell population is > 90% in CRISPR-Sirius C19q-sgRNA cells. No significant differences in growth rates were observed. NECTIN2 is located at 44.8 Mb and targeted 8 times by C19q-sgRNA. TOMM40 is located at 44.9 Mb and targeted 1 time by C19q-sgRNA. TNNT1 is located at 55.1 Mb and targeted 16 times by C19q-sgRNA. The gene locations are indicated in the diagram at the left side. C19q targeting region is marked by the red box. All experiments were repeated at least three times.

### Osteosarcoma cells have relatively high levels of CTCF, cohesin, and the euchromatin mark H3K27ac

To investigate the factors regulating large-scale chromosome conformation in osteoblasts and OS, we examined the levels of core histones and histone PTMs and the chromatin architectural proteins, which control the primary level of chromatin compaction, and mediate interphase chromatin looping, respectively. Core histones are generally responsible for ∼7-fold compaction of interphase DNA(61). Histone PTMs, especially euchromatin and heterochromatin marks, can affect chromatin folding and dynamics(62). Core histones and histone PTMs were detected by western blotting in both whole-cell lysates and acid extracts. No significant changes were found in expression levels of the core histone proteins H2A, H2B, and H3 between osteoblasts and OS cells (Figure 3B). Next, we examined the euchromatin histone PTMs H3K4me2/3 and H3K27ac, and the heterochromatin histone PTMs H3K9me2/3 and H3K27me3. While the levels of H3K4me2/3 and H3K9me2/3 were similar, we observed reduced H3K27me3 and relatively increased H3K27ac in OS cells (Figure 3C). H3K27me3 is a facultative heterochromatin mark catalyzed by PRC2 complexes. Misregulation of H3K27me3 is often found in developmental disorders and during cancer progression(63–68). We hypothesized that maintenance of H3K27me3 is required to maintain the *in vivo* chromosome conformation in osteoblasts that have high equilibrium levels of the mark but not found in OS cells.

**Figure 3.**
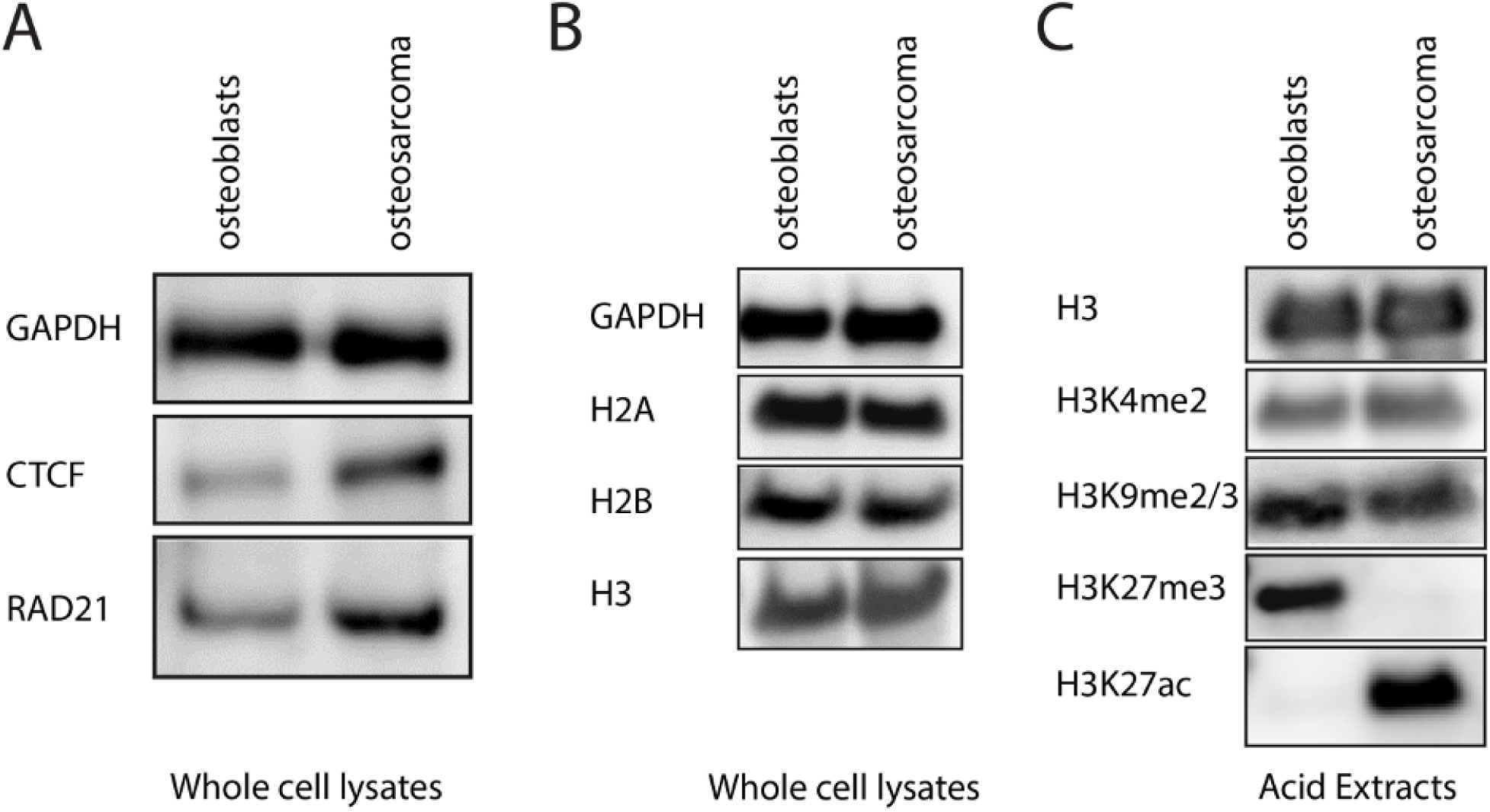
Different levels of chromatin architectural proteins and histone modifications were detected in osteoblast and osteosarcoma cells. (A) Western blotting shows that protein levels of CTCF and RAD21 (a cohesin subunit) are higher in osteosarcoma cells. (B) Western blotting of core histones. (C) Western blotting on the extraction of histone by acid extraction. While H3K9me2/3 and H3K4me2 levels are similar, a gain of H3K27ac and a loss of H3K27me3 were found in osteosarcoma. One million cells were used in each sample. All experiments were repeated at least three times.

Next, we performed western blotting for CTCF and RAD21 (the kleisin subunit of cohesin) to examine the levels of CTCF and cohesin. Depletion of RAD21 impairs cohesin function and leads to a genome-wide loss of chromatin loops(69). We found that both CTCF and cohesin are more highly expressed in OS cells than in osteoblasts (Figure 3A). We hypothesize that CTCF and cohesin promote the collapsed chromosome conformations, with increased globularity, through extensive chromatin looping. Also, we hypothesize that the extended chromatin conformation in osteoblasts is less likely regulated by cohesin/CTCF machinery whereas OS cells are more likely regulated by this machinery, leading to the collapsed/globular chromosome conformational state.

### Depletion of RAD21 alters gene expression and promotes extended chromosome conformation in osteosarcoma cells

To test whether CTCF and cohesin facilitate the formation of collapsed chromosome conformations in OS cells, we knocked down either CTCF, RAD21, or both, using RNAi. CTCF and cohesin work concomitantly to establish and stabilize chromatin loops (Figure 4A)(70, 71). The successful knockdown of CTCF and RAD21 in OS cells was demonstrated by western blotting (Figure 4B). Rhodes and colleagues reported enhanced regulation by the polycomb repressive system in the absence of cohesin in mouse embryonic stem cells (72). As a control, we examined whether the same situation occurred in OS by measuring H3K27me3 levels in siControl and siRAD21 cells (Figure 4C). We found that the H3K27me3 level remains low in siRAD21 cells, indicating that H3K27me3 is unlikely to play a dominant role in organizing C19q in the absence of RAD21 in OS.

As shown in Figure 4D, the knockdown of CTCF resulted in C19q area expansion rather than an extension of C19q conformation. This observation is consistent with the anchoring role of CTCF in the loop extrusion model. We observed a twofold increase in the C19q area in CTCF-knockdown OS cells, suggesting that continuous, deregulated chromatin looping occurred in the absence of the anchors leading to loss of linearization (Figure 4E). In accordance with these results, we observed extended C19q conformations in RAD21-knockdown cells, indicating that indeed the excessive chromatin looping indeed resulted in collapsed C19q conformations in OS cells (Figure 4D and 4F). The double knockdown of both CTCF and RAD21 also resulted in extended C19q conformations in OS cells, similar to RAD21 single knockdown (Figure 4D and 4F). The viability of OS cells 24 hours after knockdown was similar to that prior to the single knockdown of CTCF or RAD21 (Figure S3). In contrast, the double knockdown (CTCF + RAD21) reduced cell viability, indicating a potential dependence of OS cell survival on CTCF and RAD21(Figure S3).

**Figure 4.**
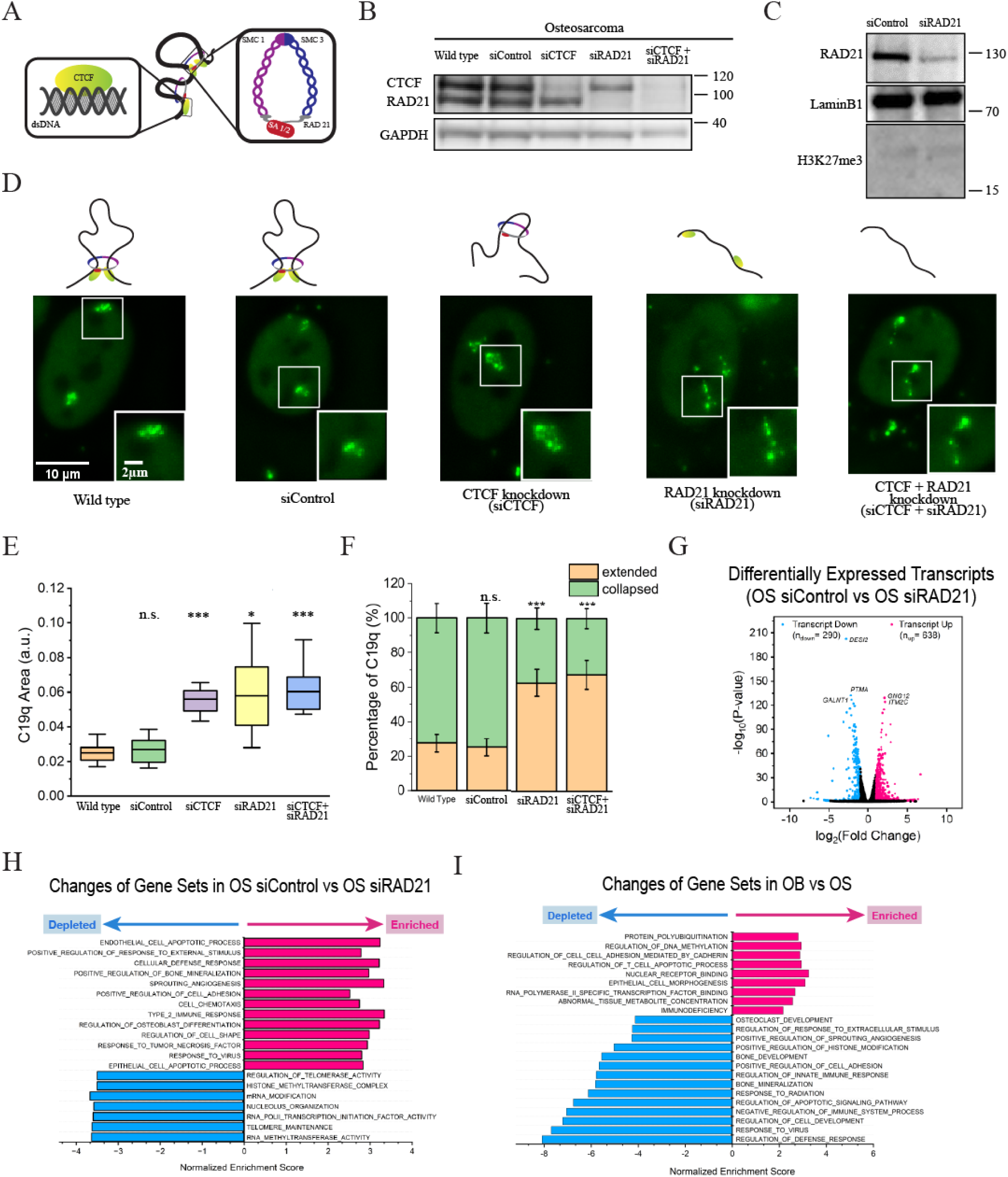
Increased chromatin looping leads to collapsed C19q conformation in osteosarcoma and different gene profile. (A) A schematic of chromatin architectural proteins, CCCTC binding factor (CTCF, green) and cohesin. RAD21 (grey) is the kleisin subunit of cohesin critical for ring formation. (B) Western blotting shows that CTCF and RAD21 protein expression levels were significantly reduced 24 hours post siRNA transfection in osteosarcoma cells (siCTCF and siRAD21 respectively). (C) Western blotting shows RAD21 knockdown does not result in changes of H3K27me3 levels in OS cells. (D) (left to right) Images of C19q in osteosarcoma cells in the indicated conditions (37°C, 5% CO_2_ and humidity). Images were taken 24 hours post-transfection for knockdown cells. (E) The area calculation of C19q in osteosarcoma cells upon siRNA transfection and negative control (siControl, N_C19q_=10 for each condition) (F) Percentages of cells with collapsed (green) and extended (orange) C19q conformations in osteosarcoma cells (N_C19q_=110 for wild type, N_C19q_=110 for negative control, N_C19q_= 110 for CTCF knockdown, N_C19q_= 110 for siRAD21, and N_C19q_= 110 for double knockdown) (G) Volcano plots of osteosarcoma cells with and without RAD21 knockdown (24 hours post-knockdown). Downregulated transcripts (log2FoldChange ≧ –1, p<0.05) are colored in blue and upregulated transcripts (log2FoldChange ≦ 1, p<0.05) are colored in pink. Top 5 genes are labeled. (H) Genome wide gene set enrichment analysis (GSEA) of RNA seq with (siRAD21) and without (siControl) RAD21 knockdown (24 hours post-knockdown, ontology, p<0.05). (I) Genome wide gene set enrichment analysis (GSEA) of RNA seq in osteoblasts and osteosarcoma cells (ontology, p<0.05). The significance tests in the figure 4E were calculated by using Welch’s T-test. The significance test in figure 4F was calculated by using Fisher’s exact test. *, p<0.05; **p<0.01, ***p<0.001; n.s., non-significant.

Next, we ask whether changes in chromosome conformation in RAD21 knockdown (siRAD21) cells are associated with changes in gene expression and cancer. We performed RNA-seq analysis of OS cells with and without RAD21 knockdown at 24 hours. Approximately 638 genes were upregulated, and 290 genes were down regulated genome-wide (absolute values of log2FoldChange equal to or greater than 1, p<0.05)(Figure 4H). GSEA in oncology showed that cancer related genes were abnormally expressed (Figure 4I). When comparing the siRAD21 GSEA to the one of OS versus OB cells (Figure 4I), we found opposite trends in bone mineralization, response to virus, positive regulation of cell adhesion, and sprouting angiogenesis genes, suggesting partial reversion of bone cancer phenotypes in OS towards the OB state in recovering immune system and cell identity. Notably, genes related to the regulation of telomerase activity, such as DKC1 and MYC, were downregulated, suggesting that RAD21 knockdown potentially reduces the lifespan of OS through telomere instability (Figure S4). These results demonstrated that changes in chromosome conformation are associated with changes in oncogenic gene expression, which is consistent with important mechanistic and clinical genomic studies on the mutations of cohesion complexes or “cohesinopathies” as tumor drivers (73).

### Osteoblasts have reduced chromatin mobility

Local chromatin stiffness can be reflected by chromatin mobility. For example, nucleolus-associated heterochromatin is less mobile than other chromatin in the nucleoplasm in HeLa cells, as observed by fluorescence correlation spectroscopy (FCS)(74). Increased levels of heterochromatin result in increased nuclear stiffness(75). To investigate the local stiffness in both osteoblasts and OS cells, we measured the mobility of the genomic locus L24, a locus with 15 copies of repeats, located within C19q at 50.1 Mb on the linear genome. We targeted the L24 locus using CRISPR-Sirius and tracked the movement of the locus over time in OS and osteoblasts (Figure 5A-5B). To quantify locus dynamics, we measured important biophysical parameters, including mean-square displacements (MSDs), trajectory radii (R_g_), and effective diffusion constants (D_eff_) (Figure 5C-5D, and S3). MSD and effective diffusion constants represent the mobility of the targeted loci. Trajectory radii are the radii of the areas being traveled by the targeted loci within a given time. We found that the L24 locus in OS is significantly more mobile and has a larger R_g_ than that measured in osteoblasts (Figure 5C-5D, and S5), indicating the relatively high stiffness of the L24 locus on chr19 in osteoblasts. Our results imply that chr19 is relatively more rigid in osteoblasts than in OS cells.

**Figure 5.**
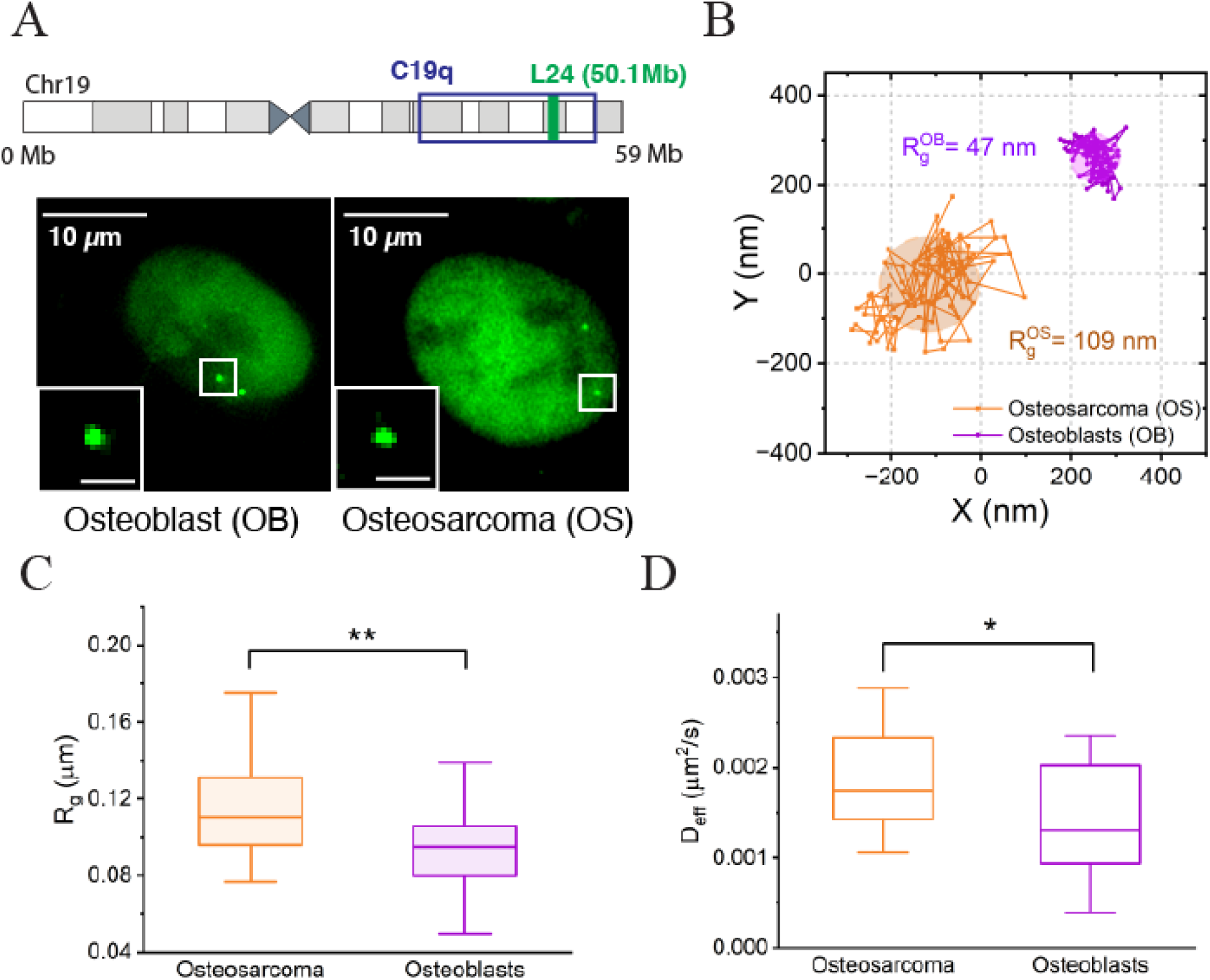
The H3K27me3-rich chromosome 19 in osteoblast cells is relatively less mobile. (A) Images of L24 loci in living cells by CRISPR-Sirius. Top: Location of L24 locus relative to the C19q targeting sites on the G-banding ideogram of human chr 19; Left: osteoblasts; Right: osteosarcoma. Images were deconvoluted (2D) to reduce the background noise. Bar in the inset, 1 μm. (B) Typical L24 trajectories in osteosarcoma and osteoblast cells. Each trajectory was obtained from tracking the locus in a video containing 100 frames with 100 ms exposure time and 0.2 sec interval. (C) Box-and-whisker plot showing the gyration radii of L24 locus trajectories (R_g_) in the osteosarcoma (n=28 trajectories, N_cell_=23) and osteoblasts (n=24 trajectories, N_cell_= 21) cells. (D) Box-and-whisker plot showing the effective diffusion constants (D_eff_) of L24 loci in the osteosarcoma (n=28 trajectories, N_cell_=23) and osteoblast (n=24 trajectories, N_cell_=21) cells. Statistical significance was assessed by unpaired Welch’s t-test with 95% confidence; \**p*<0.05 and \*\**p*<0.01. All experiments were repeated at least three times.

### H3K27me3 is essential for maintaining the extended C19q conformation in osteoblast cells

To confirm that H3K27me3 is present in the C19q region of osteoblasts and is significantly reduced in OS, we performed spike-in ChIP-seq in both cell lines. Our results confirmed that H3K27me3 is enriched within the C19q region in osteoblasts but not in the OS cells (Figure 6A). We asked whether the presence of H3K27me3 is essential for the maintenance of extended chromosomal conformations in osteoblasts. We tested this hypothesis by detecting chromosome conformations in osteoblasts with reduced H3K27me3 levels. H3K27me3 is deposited by EZH2, the catalytic subunit of polycomb repressive complex 2 (PRC2)(66, 76). To reduce H3K27me3, we used the small EZH2 inhibitor EPZ005687, which inhibits H3K27me3 deposition by binding to the SET domain of EZH2(65) (Figure 6B). H3K27me3 in osteoblasts was significantly reduced on day 8 with 5μM EPZ005687 treatment in osteoblasts, as verified by western blotting (Figure 6C). The number of collapsed C19q conformations was significantly greater in EPZ005687-treated osteoblast cells than in untreated cells whose C19q conformations are primarily extended (Figure 6D). Approximately 68% of the EPZ005687-treated cells had collapsed C19q while only 28% of the untreated cells had collapsed C19q (Figure 6E, n=50, \*\*\**p*<0.001). As control, we did not observe significant changes on the levels of RAD21, CTCF, and H3K9me3 in OB cells with reduced H3K27me3 (Figure 6F). Thus, it is unlikely that TADs or the constitutive heterochromatin mark H3K9me3 play dominant roles in collapsing C19q in reduced H3K27me3 OB cells. Our results demonstrate that H3K27me3 is essential for maintaining C19q extension the osteoblast cells, perhaps through providing necessary structural rigidity, either directly through the functional effects of the PTM on nucleosomes or through the localization of “reader” modules which can bind to this mark and are known to alter 3D chromatin conformation (5).

**Figure 6.**
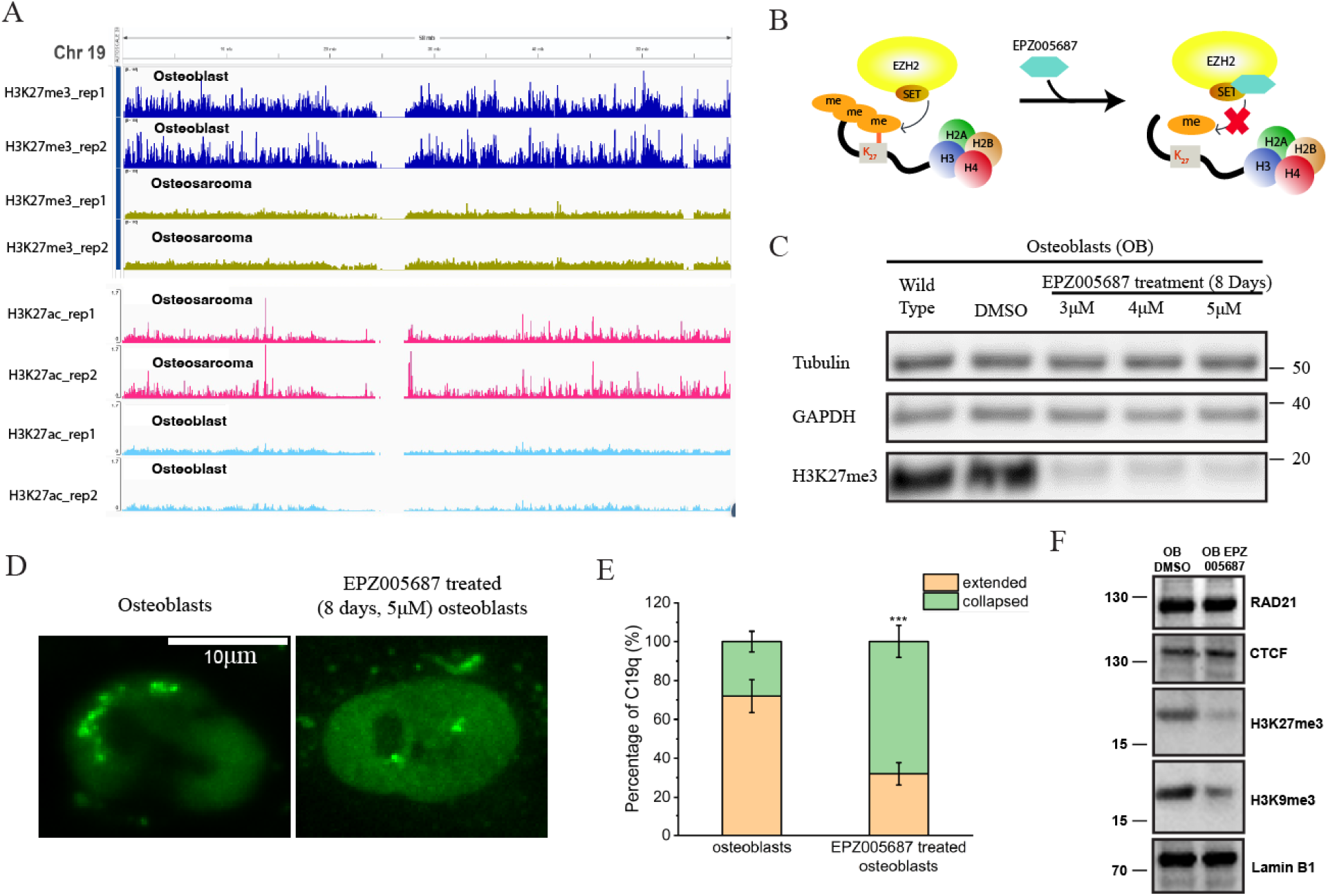
H3K27me3 is essential for maintaining extended C19q conformations in osteoblasts. (A) ChIP seq data showing H3K27me3 and H3K27ac occupancy along chr 19 in OB and OS cells. (B) Schematic diagram illustrating the action of EPZ005687, which prevents deposition of H3K27me3 by blocking the SET domain of EZH2 in PRC2 complexes. (C) Western blotting shows that H3K27me3 expression levels were significantly reduced within 8 days post EPZ005687 treatment in osteoblast cells. (D) Images of untreated osteoblast cells with extended C19q conformations and EPZ005687 treated osteoblast cells with collapsed C19q conformations. (E) Percentages of cells with collapsed (green) and extended (orange) C19q conformations in osteoblast cells under indicated conditions (N_C19q_=50 for each condition). The significance was calculated using Fisher’s exact test. \**p*<0.05, \*\**p*<0.01, ***, p<0.001. (F) Western blotting shows that reduction of H3K27me3 in OB upon EPZ005687 treatment does not change the levels of Rad21 and CTCF. However, a mild reduction of H3K9me2/3 was observed, suggesting collapsed C19q was not due to the increase of CTCF, cohesin, or H3K9me2/3.

## DISCUSSION

Interphase chromosomes have been traditionally investigated and represented as loops within DNA regions smaller than 1 Mb. Changes in 3D genome architecture at the chromatin loop scale have been reported in several diseases including cancer (24). However, opposite cases were reported that alterations of single TAD boundary had no significant effects on neighboring genes, suggesting that TADs are not the sole regulators for controlling gene expression (77). Our knowledge about the assembly, regulation, dynamics, and function of megabase-sized chromosomal domains, such as compartments, *in vivo* is still emerging. Although challenging, visualizing temporal changes in large-scale chromosome conformations in live cells is necessary to study the 3D chromatin architecture and its function, especially in diseases. In this study, we used CRISPR-Sirius, a CRISPR-based live-cell imaging technique, to monitor and track genomic loci, chromosome conformation, and dynamics in real time in OS and osteoblast cells. Due to the previously established association of chr19 with cancer development and progression(48–53), we monitored the C19q conformation (∼17 Mb). We are the first to characterize and report the extended C19q conformation in human osteoblasts as opposed to the predominantly collapsed C19q conformation in OS. Before our work, extended chromosome conformations have not been characterized in human.

Transcription has been inferred to be a “modulator” that actively shapes the 3D chromatin conformation(29, 78–80). In this scenario, transcription decompacts local chromatin conformation at the single-gene level to promote accessibility. For instance, chromatin decompaction occurs during transcriptional activation of the heat shock gene, *HSP70,* and highly expressed long genes, such as *Ttn* (29, 81). However, changes at finer-scales that code for single genes do not necessarily result in changes of chromosome territories at tens of Mb scales, which contain non-coding DNA and silenced genes in addition to actively transcribing genes. In this study, we detected no significant effects on the C19q conformation upon the inhibition of transcription using DRB, indicating that transcription alone is not sufficient to alter large-scale chromosome conformation. Similarly, the morphology of human chromosome 11 territories was unaffected by actinomycin D treatment in early G1 HT-1080 cells(60). However, in DRB-treated human primary lymphocytes, chromosome 19 territories were reported smaller than those in untreated fixed lymphocytes(82). Discrepancies among groups can potentially be explained by differences in chromatin properties modulated by DNA binding proteins and their chemical modifications in different types of cells. With CRISPR-Sirius, we showed that, at larger than ten megabass scales, the chromosome conformation is regulated by the repressive histone methylation marks H3K27me3 and chromatin architectural proteins, CTCF and cohesin complexes.

Our data demonstrate that CTCF knockdown leads to expansion rather than extension of the C19q conformation, and knockdown of RAD21 leads to an extended C19q conformation in the OS cells, likely due to loss of chromatin looping(83). Although our results align well with the loop extrusion model, contrasting evidence has been reported by other studies, indicating that chromatin architecture can be regulated by mechanisms other than TADs in different organisms (30, 84). In addition to cell type-dependent phenomena, sample preparation requiring different heat, fixation, and hybridization treatments may lead to the observations compatible and not sensitive to those reagents. Evidence for that has been reported by others and suggested by our data. For example, fixation reagents were reported to shrink nuclei due to its dehydration effects (85). Significant chromatin rearrangement was reported under temperature stress due to the repositioning of architectural proteins from TAD boundaries to interior TADs, suggesting that chromatin organization can be heat sensitive (30). We did not observe extended C19q in RAD21 knockdown cells when the samples were imaged at room temperature (∼21℃) in the absence of CO_2_ and humidity supplies (Figure S6). CRISPR-based imaging does not require insertion of foreign DNA and treatment of high temperature and hydrophobic reagents (e.g., formaldehyde), thus largely preserving the endogenous chromosome organization. Our results demonstrated the importance of the regulatory looping machinery, and also appropriate temperature and humidity conditions for 3D genome organization.

Unlike yeasts, H3K27me3 as a heterochromatin mark is involved in many cellular processes and plays a critical role in organizing the genome in higher eukaryotes (86). We reported that H3K27me3 is essential for maintaining a rigidified or extended C19q in the osteoblast cells, which has an interesting context in the literature(87). Notably, osteoblast cells have relatively low levels of cohesin and CTCF compared to those in OS (Figure 3A), implying reduced perturbations on polycomb-dependent chromosome interactions by cohesin(72). We proposed a model categorizing chromosome conformations under different conditions (Figure 7). We reasoned that the extended conformation only exists in a nuclear environment with high levels of heterochromatin marks and less structural maintenance of chromosome (SMC) complexes, such as cohesin (Figure 7, Type II). Highly rigid and collapsed conformations can be found when both SMC complexes and heterochromatin marks are enriched (Figure 7, Type I).

**Figure 7.**
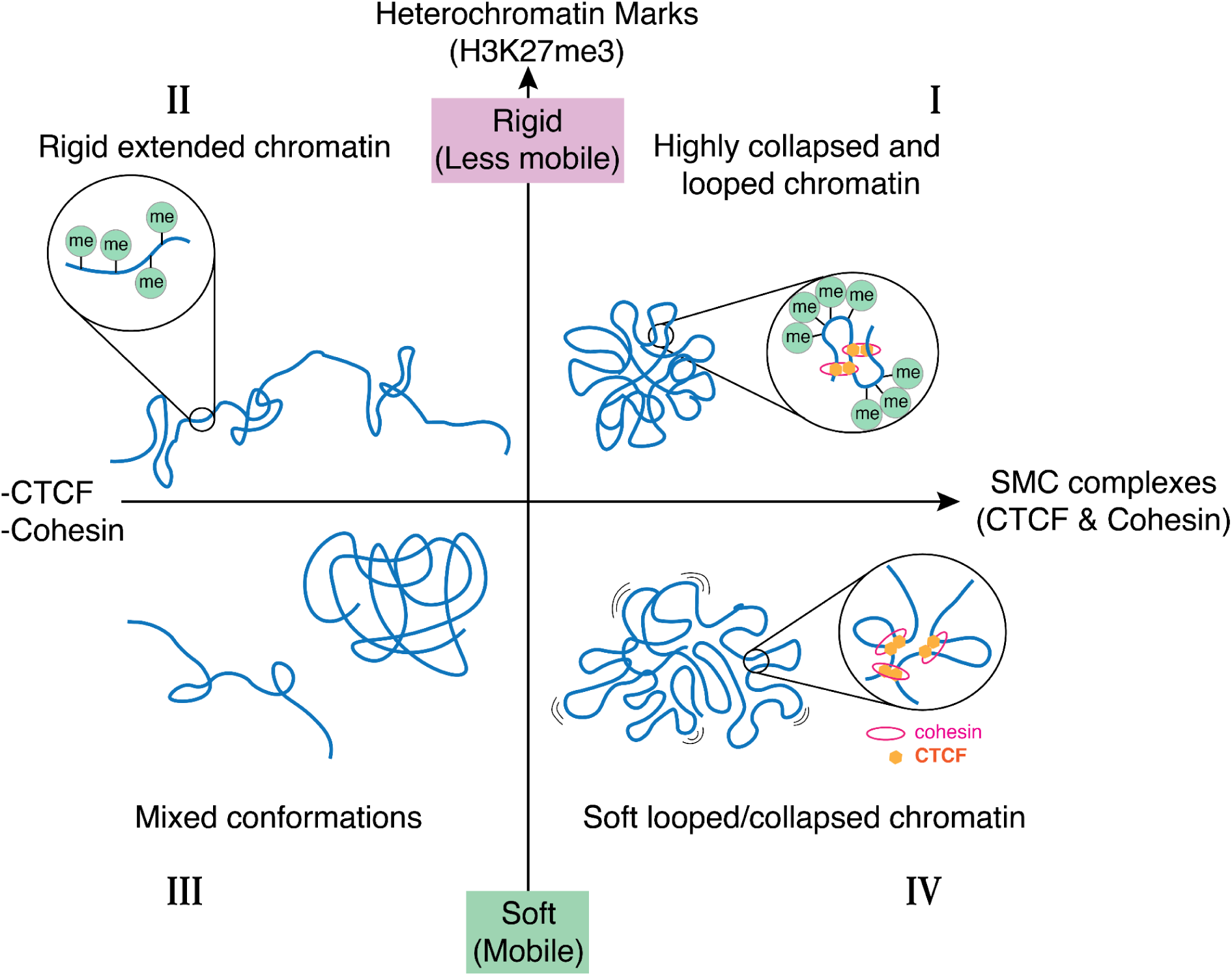
A model categorizing chromosome conformations in different conditions. Large-scale chromosome conformations are regulated by several factors. High levels of heterochromatin marks increase the rigidity of the chromatin, which results in stable conformations and slow mobility. On the other hand, the structural maintenance of chromosomes (SMC) complexes regulate the volume of large-scale chromosomal domains by chromatin looping. In type I conformation, where both heterochromatin marks and SMC complexes are abundant, highly looped and rigid chromosomes result in highly collapsed conformations with minimal volume and mobility. The type II conformation is found in the nuclear environment with fewer SMC complexes but high levels of heterochromatin marks, making chromatin less bendable and more rigid. When both heterochromatin marks and SMC complexes are low, reduced constraints on chromatin lead to relatively unstable chromosome conformations that might switch between extended and collapsed conformations. The metastable conformations are likely determined by other factors, such as intrachromosomal interactions, chromatin-organelle attachments, or transcription activities. Finally, restricted by SMC complexes but not the heterochromatin marks, individual chromosomes form a collapsed but mobile conformation found in osteosarcoma cells (Type IV).

Previous studies have shown that the presence of the heterochromatin mark H3K9me2/3 and its binding protein HP1 (heterochromatin protein 1) increases chromatin stiffness in the cells(88, 89). We characterized chromatin mobility in both osteoblasts and OS cells as an indication for chromatin rigidity. C19q in osteoblast cells is less mobile than that in OS cells, indicating that H3K27me3 likely imparts stiffness to chromatin, similar to the H3K9me2/3 mark. Two possible explanations for increased rigidity could be that (1) the presence of H3K27me3 leads to tighter packaging of nucleosomes on chromatin, which provides chromatin stiffness, thereby stabilizing the extended C19q conformations, and (2) high levels of H3K27me3 recruit chromatin reader domains, which generates steric effects, leading to reduced chromatin bendability. A combination of both scenarios likely explains the final conformation and properties of the chromosome. Future studies on chromatin rigidity are needed to further understand these mechanisms.

Simultaneously with the significant decrease in H3K27me3 in OS, we observed an increase in H3K27ac (Figure 6A). Interestingly, the increase of histone acetylation by inhibition of HDACs has no effects on the morphology of human chromosome 11 territories in HT-1080 cells (60). We speculate that the loss of H3K27me3 and increase in H3K27ac could play an important role in regulating gene expression in OS, to potentially augment oncogenic transcription. Of note, the loss of H3K27me3 at promoter regions did not always lead to the upregulation of genes, suggesting that the loss of chromatin repression is necessary but not sufficient to activate transcription(90). While our results provide exciting disease context for the loop extrusion model, and new insights into the dynamic interplay between cohesin and Polycomb functions, further work is necessary to (1) evaluate the generalizability of our model (i.e. does H3K27me3 exist in lower levels across all human OS, and is this resulting in a chromatin architectural alteration common within the tumor?) and (2) to understand the role of CBX and EED chromatin reader modules in the context of our model, whereby chromodomain containing reader proteins would impart stiffness through a distinct pathway from the electrostatics of histone PTMs.

## MATERIALS AND METHODS

### Data availability

The Raw ChIP-seq sequence files prepared for this manuscript are publicly available through the Gene Expression Omnibus (BioProject accession number: PRJNA1111443). Other data reported in this paper are available from the lead contact upon reasonable request.

### Experimental Details Plasmid construction

The sgRNA (pLH-sgRNA-Sirius-8XPP7, Cat#121940), dCas9 (pHAGE-TO-dCas9-P2A-HSA, Cat#121936), and GFP-fused PP7 coat protein (pHAGE-EFS-PCP-GFPnls, Cat#121938) expression constructs were obtained from Addgene. The design of these plasmids can be found in previous work(33, 34). The sgRNA targeting sequences, TCCCTCAGACCCnGG for C19q and AGGTGGTTAGGAnGG for L24, were purchased from Integrated DNA Technologies, and cloned into the sgRNA vector using BbsI sites. The cloned sgRNA constructs were confirmed by Sanger sequencing.

### Cell culture and transcription inhibition

Human bone osteosarcoma epithelial (U2OS), human embryonic kidney epithelial (HEK293T), and human osteoblast cell lines (hFOB 1.19) were acquired from ATCC. U2OS cells were maintained in DMEM (Corning) with high glucose supplemented with 10% FBS (Avantor) and 1% penicillin/streptomycin (Sigma). HEK293T cells were maintained in Iscove’s modified Dulbecco’s medium supplemented with 1% GlutaMAX, 10% FBS and 1% penicillin-streptomycin. The human osteoblast cell line was maintained in DMEM/F-12, 50:50 (Corning) media with L-glutamine supplemented with 10% FBS and 0.3 mg/ml G418. All the cell lines were confirmed to be free of mycoplasma contamination by using a Mycoalert PLUS kit (Lonza). To inhibit transcription, a final concentration of 50 μg/ml polymerase II elongation inhibitor 5,6-dichloro-1-β-ribofuranosylbenzimidazole (DRB; Sigma-Aldrich) was added to the culture medium and images were collected at different time points as indicated in Figure 2.

### Lentiviral transduction

To create stable cell lines with fluorescently labeled C19q, lentiviral particles carrying the sgRNA plasmids were generated using HEK293T cells as described previously(91). Approximately 5×10^6^ cells were seeded in each well of a 6 well plate 24 hours before transfection. Then, 0.5µg of pCMV-dR8.2 dvpr (Addgene), 0.3µg of pCMV-VSV-G (Addgene), and 1.5 µg of the sgRNA plasmid were cotransfected into the cell line using TransIT transfection reagent (Mirus) according to the manufacturer’s protocol. 48 posttransfection, the virus was collected and filtered through a 0.45µm filter. The virus was then immediately used or snap frozen and stored at –80°C. The hFOB 1.19 and U2OS cells maintained as mentioned above were transduced by spinfection in 6-well plates. Approximately 2×10^5^ cells were combined with 1ml of lentiviral supernatant and centrifuged at 1,200 x g for 30 minutes.

### Flow cytometry

To select dCas9 and PCP-GFP positive cells, hFOB cells were sorted by fluorescence-activated sorting (FACSAria Fusion cell sorter; BD). The FACSAria Fusion cell sorter was equipped with 405-, 488-, 561-, and 637-nm excitation lasers. PCP-GFP was detected with a 450/50 nm emission filter, and dCas9 positive cells were detected using an Alexa647-conjugated antibody to mouse CD24 (BioLegend) following the manufacturer’s staining protocol with a 670/30 nm emission filter.

### EZH2 inhibitor treatment

Human osteoblast cells (hFOB1.19) were cultured at a confluence of 60-70% and then were split back to 30% confluence in fresh medium supplemented with the small molecule EZH2 inhibitor-EPZ005687 (selleckchem.com, CAT#: S7004) on days 3 and 6. The cells were incubated with EPZ005687 at 2μM, 3μM 4μM and 5μM in a total volume of 2 ml for days 7 and 8. Western blotting was used to confirm the protein expression levels of H3K27me3.

### Fluorescence microscopy and image processing

A custom-built Olympus IX83 microscope equipped with three EMCCD cameras (Andor iXon 897), a LED, 60X apochromatic oil objective lens (NA 1.5), and mounted with a 1.6X magnification adaptor, resulting in a total magnification of 96X. The microscope incubation chamber was maintained at 37°C supplied with 5% CO2 v/v and humidity for live cell imaging. Image data were acquired using CellSens software (version 4.1.1). For C19q imaging, 10 z-slices per nucleus were acquired with a step size of 0.25µm and an exposure time of 100 ms. The images of a z-stack series were projected to a 2D image by using maximum intensity projection.

The images were processed using Fiji(92) and Mathematica (Wolfram v12.3.1). To eliminate the drift movement of cells, the movement of individual genomic loci was calibrated by the motion relative to the nuclear centroid. The mean square displacement (MSD) of lag time kΔt was calculated by (93).

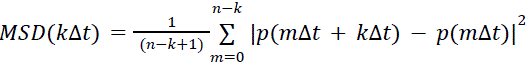

where p(t) is the position vector of a locus at time t, and Δt is a fixed time interval between two successive image frames. The gyration (or trajectory) radius R_g_ of the locus trajectory was calculated as:

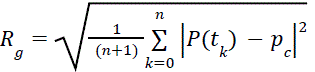

where 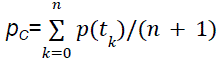 is the geometric center of the positions defining the trajectory and 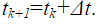. The gyration radius Rg measures the size of the area covered by locus movement. The effective diffusion constant D_eff_, is calculated by MSD at the short time intervals within 3Δt and fitted by the equation, MSD=4D_eff_ Δt(36).

The C19q conformation or shape can be characterized by asphericity which is defined by the eigenvalues of gyration tensors. The components of gyration tensor *Q* for N position vectors are defined by

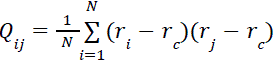

Where *r_i_* and *r_c_* denote individual denote individual position vectors and the position vector of the centroid of the N position vectors. For 2D data, *Q* is a 2×2 symmetric matrix. Let the eigenvalues of *Q* be λ_1_ and λ_2_ and λ_1_ ≤ λ_2_, asphericity Δ is defined as

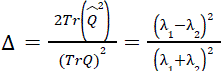

where 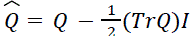 and *I* is the identity matrix (55, 56). Note that 0 ≤ Δ ≤ 1. Δ = 0 represents a circular shape while Δ = 1 represents a fully extended shape. We can also define the aspect ratio of principal axes, 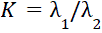. The larger K, the more extended shape. The gyration tensor *Q* of C19q was calculated from the spline curve of C19q output from Fiji (92). Using the distribution of K values, we modeled C19q conformations by Gaussian bimodal fitting using OriginLab (version 2024b).

### Whole cell protein extraction

U2OS and hFOB cells were cultured in 6-well plates, trypsinized, and collected by centrifugation at 500 g for 3 minutes. The cell pellets were washed once with PBS. A total of 1×10^6^ cells were then resuspended in 250µl of cell lysis buffer (1X RIPA, 8M urea and 1X PMSF). The samples were then sonicated with a Diagenode Bioruptor (UCD-200) at medium intensity (200 W) for 5 minutes, cycling from 30 seconds ON to 30 seconds OFF. The protein concentration was determined using a Qubit protein kit (Invitrogen, Q-33211). The protein lysates were then aliquoted and stored at –20°C until further use.

### Acid extraction of histones

To extract histones, cells were cultured in 10cm dishes, and 1×10^6^ cells were washed twice with ice-cold PBS and collected by centrifugation at 500 g for 3 minutes. The cell pellets were then resuspended in 1 ml of Triton Extraction Buffer (TEB, 0.5% v/v Triton X-100, 2mM PMSF, 0.02% w/v NaN_3_ in PBS) followed by incubation on ice for 10 minutes. The cells were collected at 375 g at 4°C for 10 minutes. Cell pellets were washed with 500µl of TEB and collected again at 375 g for 10 minutes at 4°C. The cells were then resuspended in 250µl of freshly prepared 0.2 N HCl and incubated at 4°C overnight for histone acid extraction. The samples were then centrifuged at 375 g and 4°C for 10 minutes. The supernatant was transferred to a fresh 1.5 ml tube and the protein concentration was determined using a Qubit protein kit (Invitrogen, Q-33211). The protein lysates were stored at –20°C until further use.

### Antibodies and Western Blots

50µg of whole cell lysates and 5µl of acid extracts were separated on precast Bis-Tris protein gels (Bolt Bis-Tris Plus gels) and transferred to PVDF membranes (Bio-rad). Rabbit polyclonal antibodies against CTCF (Cat#2899S), H2B-V119 (Cat#8135S), H3 (Cat#9715S) and rabbit monoclonal antibody against H2A-D603A (Cat#12349S) were obtained from Cell Signaling Technology. Rabbit polyclonal antibodies against RAD21 (Cat#27071-1-AP), GAPDH (Cat#10494-1-AP) and alpha-Tubulin (Cat#11224-1-AP) were obtained from Proteintech. Rabbit polyclonal antibody for H3K9me2/3 (Cat#ab8898) was obtained from Abcam. Rabbit polyclonal antibody against H3K27me3 (Cat#39155) was obtained from Active Motif.

### siRNA transfection

U2OS cells were seeded at 6×10^4^ cells per well in a 24 well glass-bottom plate or at 2×10^4^ cells in the microscope imaging dish. The siRNAs against CTCF (ID: s20966, cat:4392420), RAD21 (ID: s531212, cat:4392420) and negative control #1(cat:4390843) were obtained from Ambion. The cells were then transfected upon reaching 60% confluency using TransIT transfection reagents (Mirus). The images were taken 24 hours post-transfection.

### RT-PCR

Total RNA was extracted by using a RNeasy plus mini kit (Qiagen), then first-strand cDNA was synthesized by using the SuperScript IV reverse transcriptase (Invitrogen) according to the supplier’s instructions. β-ACTIN was used as an internal control for RT-PCR analysis. DNA was stained by ethidium bromide. DNA oligonucleotides used in this study: NECTIN2-F: GGAAACCGGAGCCCAGGATG, NECTIN2-R: GAGACGAAGGACAGCCGCTC, TOMM40-F: GAGGAGTGCCACCGGAAGTG, TOMM40-R: CCCACCAGTACAGGGAACGC, TNNT1-F: AGGAGGAAGCCCCCGAAGAG, TNNT1-R: CCTCCTTCTTCCGCTGCTCG, β-ACTIN-F: GTGGGGCGCCCCAGGCACCA, and β-ACTIN-R: CTCCTTAATGTCACGCACGATTTC.

### RNA seq analysis

The raw data of RNA-seq for RAD21 knockdown (GEO accession no. GSE89799, SRA – SRX2344649, SRX2344650, SRX2344659, and SRX2344660) and hFob1.19 (GEO accession no. GSE118488, SRA – SRX4549277, SRX4549278) and untreated U2OS (GEO accession no. GSE118488, SRA – SRX4549306 and SRX4549307) were obtained from Sequence Research Archive (94). The data analysis, including DESeq and GSEA, was performed on SciDAP (https://scidap.com/).

### Chromatin immunoprecipitation

The U2OS and hFOB cells were cultured in 10 cm dishes and trypsinized. About 3.15×10^6^ cells were collected by centrifugation at 500 g for 5 minutes and then washed with PBS. The cell pellets were resuspended in 10 ml fixing buffer (50mM Hepes pH 8.0, 1mM EDTA, 0.5mM EGTA, 100mM NaCl). 16% w/v methanol-free formaldehyde (thermo-scientific, Cat#28906) was added to a final concentration of 1% for cell fixation. After an incubation of 10 minutes at room temperature the reaction was quenched using glycine at a final concentration of 125mM. The cell suspension was placed on ice for 5 minutes followed by centrifugation at 1200 g at 4°C for 5 minutes and the supernatant was discarded. The 2×10^6^ formaldehyde fixed mouse C2C12 myoblasts were spiked in with each sample containing 3.15×10^6^ human cells resuspended in NP1 Rinse 1 buffer (50mM Hepes pH 8.0, 140mM NaCl, 1 mM EDTA, 10% glycerol, 0.5% NP-40, 0.25% Triton X-100). Samples were incubated for 10 minutes at room temperature and pelleted at 1200 g for 5 minutes at 4°C. The cell pellets were resuspended in NP Rinse 2 buffer (10mM Tris-Cl pH 8.0, 1mM EDTA, 0.5mM EGTA, 200mM NaCl) followed by centrifugation at 1200 g for 5 minutes at 4°C. Finally, the cells were washed with Covaris Shearing buffer to remove any salts. The cells were then resuspended in 900ul of Covaris shearing buffer after adding freshly prepared protease inhibitor cocktail (Sigma, Cat# 11697498001) to the Covaris shearing buffer. The cells were then transferred to 1 ml Covaris shearing tube (Covaris milliTUBE 1ml AFA Fiber, 520135). The cells were sonicated using Covaris sonicator (E220 evolution) for 8 minutes at a cycle of 30 seconds ON and 30 seconds OFF. 12.5µl of the sonicated chromatin stock was added to 12.5µl of Covaris shearing buffer along with 10µg RNase A (Thermo Scientific, EN0531) and incubated at 37°C for 30 minutes. 10µg of proteinase K (Zymo Research, D3001-2-20) was added to this input sample followed by an overnight incubation at 65°C. The input sample was then purified using MinElute PCR purification columns (Qiagen, 28004). To check the shearing, 5µl of eluted DNA was run on E-gel 2% EX agarose gel (invitrogen, G401002) for 10 minutes. The remaining 960µl sheared chromatin lysate was centrifuged at 10,000 g at 4°C for 5 minutes. Combine the supernatant with 240µl of 5xIP buffer (250mM HEPES/KOH pH 7.5, 1.5 M NaCl, 5mM EDTA, 5% Triton X 100, 0.5% Sodium Deoxycholate, 0.5% SDS). 5µl of H3K27me3 antibody (Active motif, Cat#39155) was added to each sample. For each sample, 40µl of Protein G magnetic Dynabeads were buffer exchanged with ChIP buffer (50mM HEPES/KOH pH 7.5, 300mM NaCl, 1mM EDTA, 1% Triton-X 100, 0.1% sodium deoxycholate, 0.1% SDS) added to the sample after the antibody was added. The antibody: chromatin mixture was incubated overnight at 4°C with overhead rotation. The beads were then washed twice with ChIP buffer, once with DOC buffer (10mM Tris-Cl pH 8.0, 0.25M LiCl, 0.5% NP40, 0.5% sodium deoxycholate, 1mM EDTA) and once with TE buffer pH 7.4. The beads were resuspended in 100µl TE buffer pH 7.4 with 2.5µl 10% SDS and 5µl 10 mg/ml Proteinase K. the ChIP samples were then reverse crosslinked at 65°C overnight. The supernatant was purified with Qiagen MinElute PCR purification columns. The ChIP was validated using real-time PCR with primers designed against HOXB7 (forward – 5’-GAGTAACTTCCGGATCTACCC-3’, reverse – 5’-CGTCAGGTAGCGATTGTAGTG-3’ known to be enriched with H3K27me3 and GAPDH (forward – 5’-ACATCGCTCAGACACCATG-3’, reverse – 5’-TGTAGTTGAGGTCAATGAAGGG-3’ as a negative control. The primers were purchased from IDT. For validation, 1µl of the input sample or ChIP sample along with validation primers and SYBR green master mix (Biorad, 1725270) was added in the BioRad RT-PCR plates. The reaction mix was incubated in Biorad CFX96 Real time system with C1000 thermal cycler module at standard RT-PCR conditions(95). Ct values from both the negative and positive control were used to calculate fold enrichment using the ΔΔCt method(95).

### ChIP library construction and next generation sequencing

ChIP and input DNA from both U2OS and hFOB 1.19 were prepared for sequencing as directed by Lucigen End-It DNA repair Kit, followed by 3’ A-tailing by Klenow exo (NEB, M0212L) and adaptor ligation using T4 DNA ligase (NEB, M0202L). The linker ligated DNA was size selected on an E-gel 2% EX agarose gel. The libraries were amplified using barcoded primers followed by gel purification to remove primer dimers or unreacted primers by gel purification. Pooled libraries were validated using Qubit and sequenced at the Nationwide Children’s Hospital Institute for Genomic Medicine (NCHIGM) on a HiSeq4000 sequencer to obtain 150bp paired end reads.

### ChIP-seq data analysis

Based on the ratio of spike-in mouse cells and human cells, we performed read-depth correction by normalizing all input and ChIP samples using the mouse-spike-in reads. The raw fastq files obtained from NCHIGM were checked for quality using the FastQC v.0.72(96). The fastq reads were aligned separately to the mouse genome build mm10 (GRCm38) and the human genome build hg38 (GRCh38) using the bioinformatic tool Bowtie2 v.2.4.2(97), to generate the BAM files. To calculate the scaling factor (f) for normalizing the samples, the minimum number of mice aligned reads (m) were identified across all samples. The scaling factor was calculated by dividing the minimum number of mice aligned reads by the number of aligned mice reads for each sample (s) – f = m/s. Samtools View(98) –s –b was used to subsample the hg38 aligned BAMs as it retains the read pair information. The subsampled BAM files were then converted to paired end fastq files using the Picard SamToFastq v.2.18.2(99).

The normalized subsampled reads were aligned to hg38 using Bowtie2 v.2.4.2(97) with parameters bowtie2 –X2000. BAM sort was used to sort the BAM files. The list of blacklist regions compiled by ENCODE consortia – hg38-blacklist.v2.bed.gz was used to remove blacklist regions(100). Similarly, unmapped, mate unpaired, multi-mapped, duplicate reads, PCR duplicates and low mapping quality reads (MAPQ > 30) were removed using ngsutils filter v.0.5.9(101) and bamtools filter v.2.4.0(102). MACS2 was used to call peaks on the filtered data using the following parameters MACS2 –p 1e-2 –nomodel –shift 0 –extsize $[FRAGLEN] –keep-dup all –broad –SPMR, where FRAGLEN is the estimated fragment length based on your library size(103). For visualization, bedGraph files were generated with MACS2 bdgcmp from the pile-up, and then converted to bigwig format using BedGraphtobiWig and visualized using Integrative Genomics Viewer (IGV)(104).

### Quantification and Statistical Analysis

All analytical and statistical tests other than ChIP-seq and RNA-seq were performed using R (4.0.5)(105).

## SUPPLEMENTAL INFORMATION

Provided as a separate file.

## ACKNOWLEDGEMENTS

We thank Dr. Mark Parthun and Dr. Prabhakaran Nagarajan, Department of Biological Chemistry and Pharmacology, Ohio State University, Columbus, Ohio, for guidance in sonication using the Covaris machine and histone acid extraction protocols. The authors also thank Dr. Jyan-Chyun Jang, Center for Applied Plant Science, The Ohio State University, Columbus, Ohio for sharing instruments for Western Blots. We thank Jenna Thuma for providing the mathematical code for C19q area estimation and Dilshodbek Nishonov for obtaining the RAD21 knockdown RNA-seq data. The authors also acknowledge the Tu lab members for their insightful discussions and comments. This study is supported by the NIH grants R00 GM126810 and R35 GR132998, and OSU start-up fund to L-C.T. Research reported in this publication was supported by The Ohio State University Comprehensive Cancer Center and the National Institutes of Health under grant number P30 CA016058.

## AUTHOR CONTRIBUTIONS

Conceptualization, MB, YCC and LCT; methodology, MB, YCC and LCT; formal analysis, YCC contributed to biophysical analysis and conformation quantitative analysis. LCT and MB contributed to the genome and transcriptome sequencing analysis; Investigation, YCC, SLH, LCT, SW, and MB performed experiments. Resources, BDS, MW, and BZS; writing original draft, MB, YCC and LCT, manuscript reviewing and editing, all authors contributed; funding acquisition and project supervision, LCT.

