## Supplemental material for "Differential Regulation of Large-scale Chromosome Conformations in Osteoblasts and Osteosarcoma"

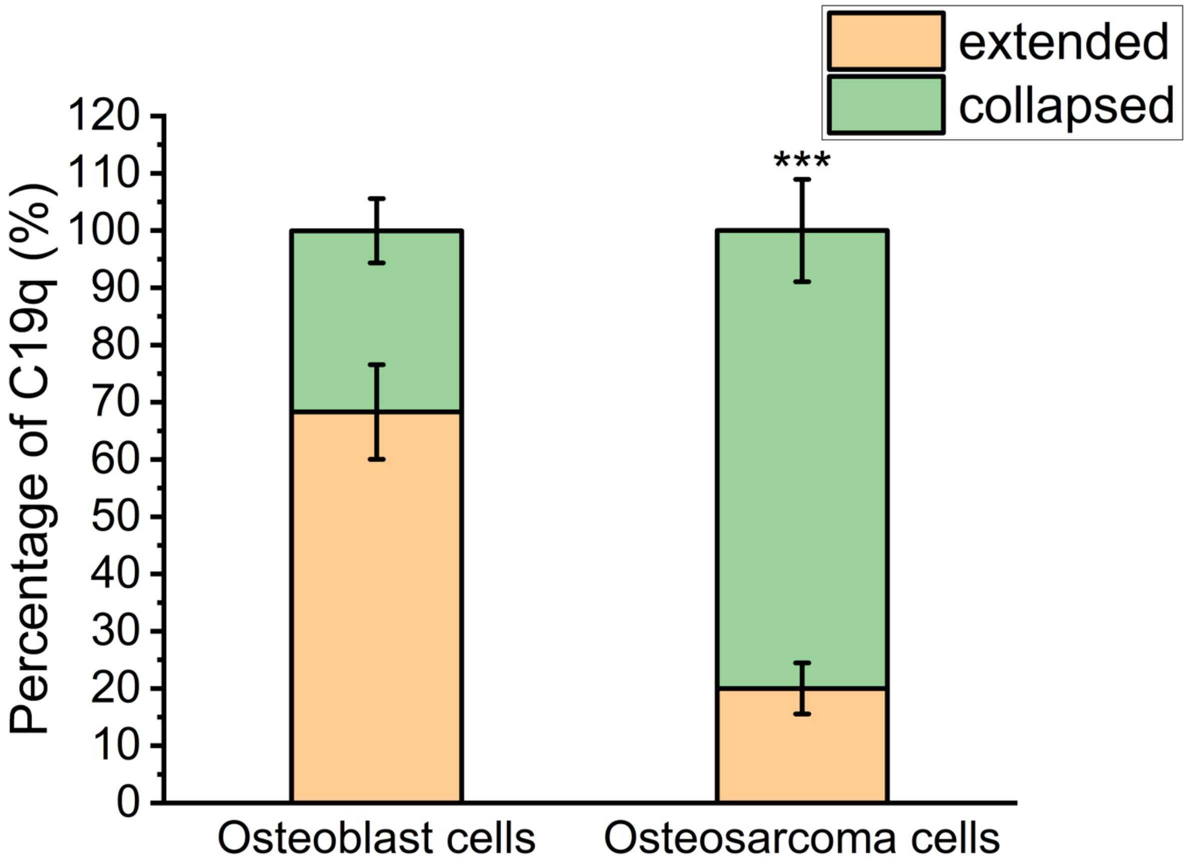


**Figure S1. Distinct chromosome conformations in synchronized osteosarcoma cells.** Percentages of cells with collapsed (green) and extended (orange) C19q conformations in osteosarcoma cells synchronized to mid-late G1 phase ($N_{C19q}$=60 for each sample). The significance test was calculated using Fisher's exact test. ***, $p<0.001$.

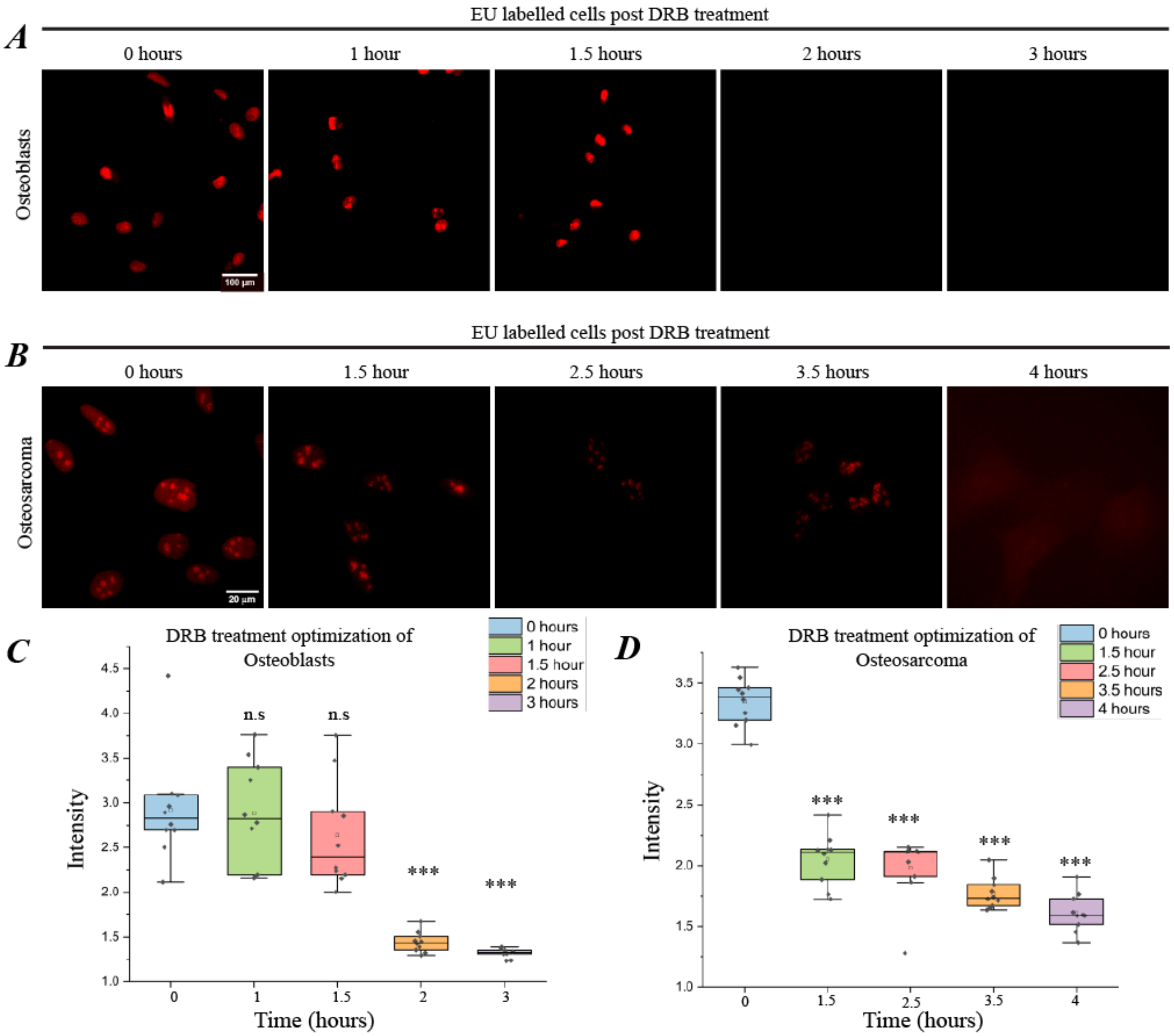


**Figure S2. DRB treatment significantly reduced newly synthesized RNA in osteoblasts and osteosarcoma cells.**

5-ethynyl uridine (EU) was used to detect newly synthesized RNA during transcription inhibition using DRB in (A) osteoblast cells and (B) osteosarcoma cells. Prolonged EU incubation induced extensive cell death. Intensity measurement of the EU signal in (C) osteoblasts and (D) osteosarcoma cells ($N_{C19q}$ =10 for each condition). The significance was calculated using Welch's T-test: significant difference, *, p<0.05; **, p<0.05, ***, p<0.001; N.S., non-significant.

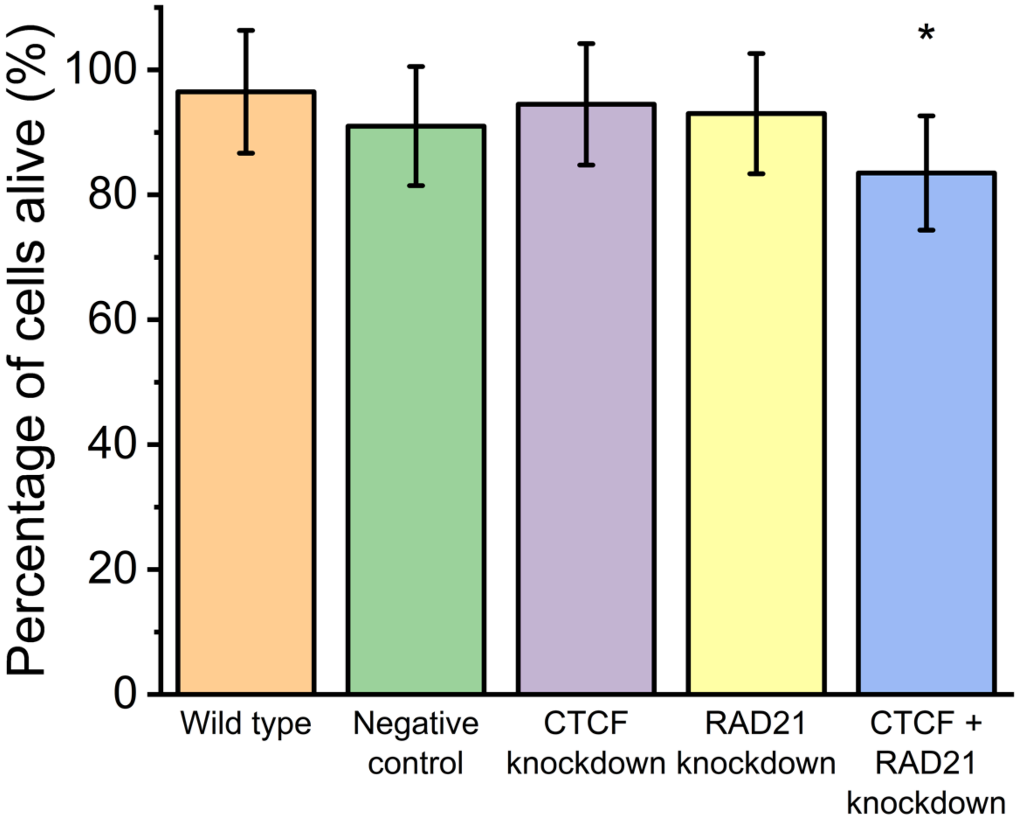


**Figure S3: Cell viability in osteosarcoma upon RNAi**
Plot shows that osteosarcoma cells have no significant change in viability upon RNAi with a negative control, CTCF and RAD21 knockdown for 24 hours post-transfection. However, the cell viability decreases in osteosarcoma cells upon double knockdown of both CTCF and RAD21(*, p-value < 0.05). P-values in the figure are calculated by using Chi-square test with Yates correction.

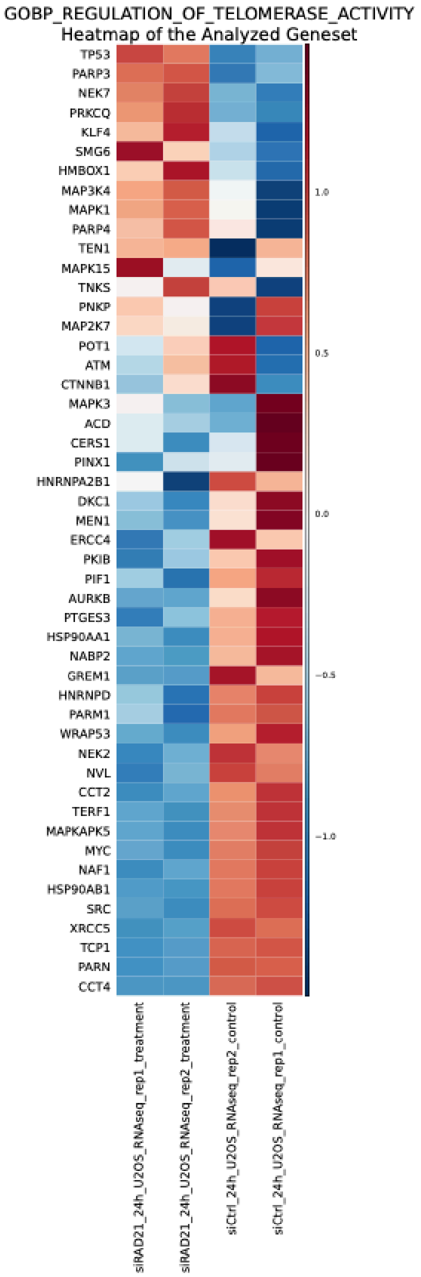


**Figure S4: Heatmap of dysregulated genes associated with regulation of telomerase activity upon RAD21 knockdown in osteosarcoma cells**

GSEA analysis in ontology showed that genes associated with regulation of telomerase activity were largely depleted upon RAD21 knockdown at 24 hours in osteosarcoma cells. Upregulated genes are colored in red and downregulated genes are colored in blue.

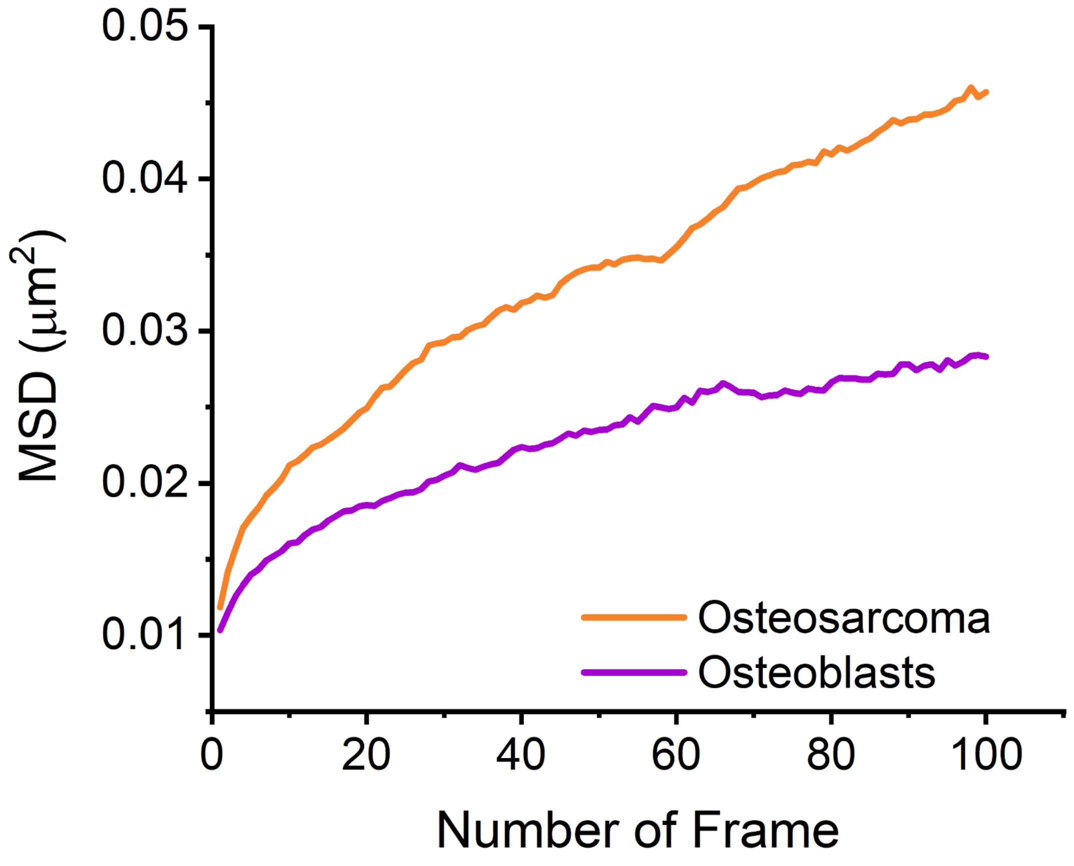


**Figure S5. MSD Curve of locus dynamics.** MSD curves of L24 genomic loci (Number of frames = 100, n = 28 trajectories for osteosarcoma, 24 trajectories for normal osteoblasts, $N_{cell} \geq 21$) in normal osteoblasts (purple) and osteosarcoma (orange) cells.

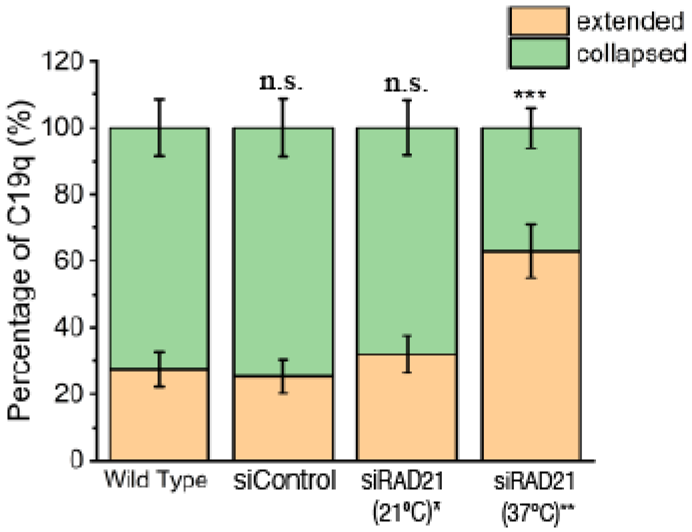


**Figure S6: C19q chromosome conformations are sensitive to cell survival conditions.** Chromosome conformations changed based on exposure to different conditions in the absence of RAD21. Percentages of cells with collapsed (green) and extended (orange) C19q conformations in osteosarcoma cells under indicated conditions (NC19q= 110 for each condition). Wild Type (no treatment) and siControl cells were imaged under the same condition as siRAD21 (37℃ with humidity and $CO_2$ supplies.), whereas siRAD21 was imaged under room temperature (37℃) without humidity and $CO_2$ supplies. The significance test was calculated by using Fisher's exact test. *, p<0.05; **p<0.01, ***p<0.001; n.s., non-significant.

**Movie S1.** This movie shows a typical movement of the C19q in the osteoblast (hFob1.19) cell nucleus recorded over 70 minutes (each 2D image is a projection from 10 z-slices). The C19q is labeled by GFP. A false green color was added for a better visualization. The movie play rate is 3 Hz. Individual images in this movie are shown in Figure 1D.

**Movie S2.** This movie shows a typical movement of the C19q in the osteosarcoma (U2OS) cell nucleus recorded over 70 minutes (each 2D image is a projection from 10 z-slices). The C19q is labeled by GFP. A false green color was added for a better visualization. The movie play rate is 3 Hz. Individual images in this movie are shown in Figure 1E.
